# Impact of high-fat Western diet on chronic lymphocytic leukemia disease progression and gut microbiome profile in Eµ-TCL1 mice

**DOI:** 10.64898/2026.03.30.715124

**Authors:** Sydney A Skupa, Jordan B Hernandez, Audrey L Smith, Erin M Drengler, Anand K Seth, Shesh N Rai, Jonathan B Clayton, Christopher R D’Angelo, Dalia El-Gamal

**Affiliations:** Eppley Institute for Research in Cancer and Allied Diseases, University of Nebraska Medical Center, Omaha, NE, 68198, USA; Department of Biology, University of Nebraska Omaha, Omaha, NE, 68182, USA; Nebraska Food for Health Center, University of Nebraska-Lincoln, Lincoln, NE, 68588, USA; Department of Genetics, Cell Biology and Anatomy, University of Nebraska Medical Center, Omaha, NE, 68198, USA; Biostatistics and Informatics Shared Resource, University of Cincinnati Cancer Center, Cincinnati, Ohio, 45267, USA; Cancer Data Science Center, College of Medicine, University of Cincinnati, Cincinnati, Ohio, 45267, USA; Department of Biostatistics, Health Informatics and Data Sciences, College of Medicine, University of Cincinnati, Cincinnati, Ohio, 45267, USA; Department of Pathology and Microbiology, University of Nebraska Medical Center, Omaha, NE, 68198, USA; Department of Food Science and Technology, University of Nebraska-Lincoln, Lincoln, NE, 68588, USA; Fred and Pamela Buffett Cancer Center, University of Nebraska Medical Center, Omaha, NE, 68198, USA; Division of Hematology, University of Nebraska Medical Center, Omaha, NE, 68198, USA; Division of Hematology and Oncology, Department of Internal Medicine, College of Medicine, University of Cincinnati, Cincinnati, OH, 45267, USA

**Keywords:** diet modulation, high-fat Western diet, gut microbiome, B-cell chronic lymphocytic leukemia, Eµ-TCL1 murine model

## Abstract

**Background:** The composition and function of the gut microbiome have been shown to contribute to both health and disease. One of the most powerful modulators of microbial composition and function is diet.

**Materials & Methods:** Using the Eµ-TCL1 murine model of B-cell chronic lymphocytic leukemia (CLL), we assigned male and female mice to a high-fat, high-carbohydrate Western diet (HF) or standard chow (CH) diet.

**Results:** Mice consuming a HF diet had significantly shorter survival than those consuming a CH diet, irrespective of sex, with female mice exhibiting particularly poor outcomes. We also observed a significant increase in splenic involvement by CLL in the HF diet-fed mice at time of sacrifice. Mice receiving the HF diet demonstrated immediate and profound effects on the gut microbiome, marked by reduced alpha diversity and significantly different community composition as measured by beta diversity. Notably, there was a sustained increase in *Akkermansia muciniphila* and *Bacteroidetes thetaiotaomicron* in HF diet-fed mice, coupled with a corresponding increase in microbiome functional pathways related to arginine and histidine biosynthesis, chitin degradation, and nucleotide biosynthesis.

**Discussion:** Collectively our data provides evidence of the profound and sustained impact of a high-fat Western diet upon the gut microbiome community and CLL pathogenesis in the Eµ-TCL1 murine model of CLL.

## 1 Introduction

The gut microbiome, a community of microbes existing within the gastro-intestinal (GI) tract of animal hosts, has been increasingly recognized for its relationship with host immunity, metabolic function, and potential contribution to the pathogenesis of oncologic diseases (1). However, there remains a significant need to understand key mechanisms driving the relationships between the gut microbiome and oncogenesis. Gut microbial dysbiosis is a potentially pathogenic gut microbiome state characterized by reductions in community diversity, presence of inflammatory bacteria taxa, and reduction of beneficial metabolite production. Disease progression in chronic lymphocytic leukemia (CLL), a mature B-cell malignancy, is associated with the development of gut dysbiosis in both animal models and human subjects (2–4). In the Eµ-TCL1 murine model of CLL, antibiotic administration caused significant changes to the gut microbiome and delayed CLL progression (2), highlighting the potential influence that regulation of the gut microbiome may have in CLL pathogenesis.

Long-term dietary intake influences the composition and functional capacity of the gut microbiome (5). Diet may also impact host homeostasis via gut microbial and non-microbial mechanisms to regulate host immune function, metabolism, and age-related diseases (6,7). Additionally, diet-induced shifts in microbial equilibrium can contribute to gut dysbiosis, oncogenesis, metabolic impairment, and many other inflammatory diseases (8–13). Studies in gnotobiotic mice colonized with human fecal microbiota demonstrated that, when switched from a low-fat, plant-polysaccharide rich diet to a high-fat, high-sugar diet, a clear shift in microbial species and metagenome composition was evident as early as 24 hours (14). Additionally, a new, stabilized microbial community was established in seven days (14). Dietary factors also undergo metabolism by gut microbes, producing an array of metabolites which play important roles in host metabolism, inflammatory signaling, immune cell function, and intestinal permeability (15). Furthermore, high-fat diets and meat-focused protein diets are associated with fewer Bifidobacteria and decreased butyrate oxidation by colonic epithelial cells, subsequently increasing intestinal inflammation and decreasing barrier integrity (16,17). High-fat diets can also raise fecal secondary bile acid levels, particularly deoxycholic acid and lithocholic acid, which are implicated in colonic cell proliferation, colonic inflammation, and subsequently colon cancer (18,19). Additionally, secondary bile acids (e.g., ursodeoxycholic acid, 7-ketolithocholic acid, hyodeoxycholic acid, murideoxycholic acid, and isodeoxycholic acid) were shown to be elevated in CLL patient plasma, specifically in patients with higher disease burden (19). These secondary bile acids found to be differentially abundant in CLL patients also displayed differential effects on T-cell function (i.e., impaired CD8+ T-cell proliferation) (19). High-fiber diets generally produce short-chain fatty acids (SCFAs) such as butyrate. Administration of an inulin (a type of prebiotic fiber) effectively modulated the gut microbiome, induced systemic memory T-cell responses, and enhanced the efficacy of anti-PD-1 therapies in an adenocarcinoma murine model (20). In lymphoma, Wei *et al.* demonstrated that a high-fiber diet increased microbial butyrate production in the blood and at tumor sites (21). The increase in butyrate elicited protective effects by exhibiting anti-proliferative and pro-apoptotic activity in both mouse and human lymphoma cells (21).

While knowledge of the role diet plays in balancing the gut microbiome in malignancies is expanding, the relationship between B-cell malignancies and dietary interventions remains poorly understood, limited to epidemiological studies with mixed results (22–24). We hypothesized that a diet more closely reflective of the average American diet – enriched in carbohydrates, fats, and low in fiber – would negatively impact the gut microbiome and influence CLL outcomes. To test this hypothesis, we performed a diet modulation study to elucidate its role in cancer initiation and progression in a murine model of CLL.

## 2 Materials and methods

### 2.1 Microbiome studies in murine models of CLL

All animal experiments were approved by the University of Nebraska Medical Center (UNMC; Omaha, NE) Institutional Animal Care and Use Committee (IACUC). All mice were housed in a specific-pathogen free (SPF) vivarium and given water ad libitum. Murine diets varied and are detailed below.

#### Diet modulation model

To investigate the effect diet has on CLL disease, a diet modulation model was utilized. In brief, two experimental diets were selected and administered to female (n = 20) and male (n = 20) mice. Experimental diets, standard chow (chow; CH) and a version of the Total Western Diet (high-fat, HF), were formulated by Teklad/Envigo and obtained as a single lot from the vendor. The HF diet is a formulated diet that represents the typical American nutrition (25). Detailed descriptions of both diets can be found in **Supplementary Table 1**. Two weeks before CLL engraftment, female and male mice (C57BL/6J) were randomized into diet groups: female chow (F-CH; n = 10), male chow (M-CH; n = 10), female high-fat (F-HF; n = 10), and male high-fat (M-HF; n = 10). Once detailed into each diet and to ensure mice were eating their respective diets, food was routinely weighed and replaced with fresh food every ten days. Correspondingly, mice were weighed every ten days to ensure proper food intake as well as to monitor any potential disease side effects (i.e., weight loss, splenomegaly) (**Supplementary Fig. S1**). Additionally, the volume of water consumed per cage was measured and replenished every ten days (**Supplementary Fig. S1**).

Two weeks after diet shift, mice (n = 40) were adoptively transferred with 0.5e^7^ Eµ-TCL1 spleen-derived lymphocytes from a moribund Eµ-TCL1 mouse (i.e., evident splenomegaly, ≥ 90% CD45+/CD19+/CD5+ cells in the blood and spleen) via tail vein injection to model CLL disease as previously described (2,26,27). A baseline (pre-CLL) blood and fecal sample was collected two-weeks post-diet shift, but prior to disease initiation. Subsequent fecal samples were collected every three weeks. Leukemic disease burden and additional lymphoid populations were evaluated via flow cytometry analysis. Due to the differential overall survival of female and male mice, female flow cytometry analyses are documented at pre-CLL, 2-weeks post-engraftment, and 4-weeks post-engraftment, while male flow analyses are documented at pre-CLL, 4-weeks post-engraftment, 8-weeks post-engraftment, and 12-weeks post-engraftment.

Mice were monitored for survival until they reached predefined humane endpoints, which included advanced tumor burden (e.g., splenomegaly or ≥ 90% leukemic peripheral blood lymphocytes), ≥ 20% weight loss from baseline, persistent hunched posture, lethargy, respiratory distress, or signs of unrelieved pain or distress. Once deemed to be moribund, mice were euthanized via CO_2_ inhalation (gradual flow rate of 30 – 70%) and cervical dislocation. Tissues (blood, spleen) were collected for further analysis. Processing details are noted below.

### 2.2 Murine sample processing

Murine blood immunophenotyping for routine evaluation of disease status (%CD45+/CD19+/CD5+ cells) and lymphoid cell populations was performed using ∼25 µL of blood obtained from the submandibular vein. Spleens, collected into sterile phosphate buffered saline (PBS) containing 2% heat inactivated fetal bovine serum (hi-FBS) (Avantor®; Radnor, PA, USA), were homogenized into a single cell suspension by passing through a 70-µm filter, then red blood cells were lysed with 1X red blood cell (RBC) Lysis Buffer (BioLegend; San Diego, CA, USA). Freshly isolated murine splenocytes were cryopreserved in 90% hi-FBS containing 10% dimethyl sulfoxide (DMSO) for further analysis.

### 2.3 Fecal microbiome sampling

Fecal pellets were collected from mice by bag collection. Pellets were collected in the morning hours, between approximately 05:00 and 09:30. Mice were placed into individual, sterile, autoclaved paper bags and left for approximately 20 – 30 minutes. Following this extended time, fecal samples were collected from each bag and aseptically transferred into sterile 1.5 mL Eppendorf tubes. The number of fecal pellets was documented at the time of collection. Fecal samples were then mechanically homogenized using autoclaved Puritan® Wooden Applicator/Stirring Sticks (Puritan Medical Products; Guilford, ME) and immediately placed on dry ice. All samples were stored in a freezer at −80°C prior to DNA extraction.

### 2.4 Fecal DNA extraction

Total bacterial genomic DNA was purified from fecal samples using the QIAamp® PowerFecal® Pro DNA Kit (QIAGEN; Hilden, Germany) and TissueLyser LT (QIAGEN) following manufacturer instructions. The quantity and quality of extracted DNA was assessed using the NanoDrop One Microvolume UV-Vis Spectrophotometer (ThermoFisher Scientific; Waltham, MA). Bacterial genomic DNA was stored at −20°C and held for no longer than one week prior to submission for shotgun metagenomic sequencing (UNMC Genomic Core).

### 2.5 Metagenomic shotgun sequencing and analysis

#### Processing pipeline

Bioinformatic processing of shotgun sequence data was performed with a custom Snakemake pipeline. After trimming adaptors with Cutadapt (minimum-length=75), reads were aligned to the host genome (GCA_000001635.9) with Bowtie2. Following removal of host reads, microbial taxonomy was classified for remaining sequences with MetaPhlAn4 (database version mpa_vJun23_CHOCOPhlAnSGB_202403). MetaCyc functional pathways were annotated with HUMAnN 3.9 using the ChocoPhlAn and UniRef90 databases (version 201901b), and pathway abundance was converted to counts per million with the renorm_table function.

#### Gut microflora analysis

Statistical analysis of shotgun data was performed by running R code in a Jupyter Lab notebook. MetaPhlAn taxa counts were converted to relative abundance with the *microbiome* package. The *phyloseq* package was used to calculate alpha diversity (Shannon index) and beta diversity (Bray-Curtis dissimilarity) at the species level. Beta diversity significance was calculated by performing PERMANOVA with marginal effects (999 permutations) using the *vegan* adonis2 function. Alpha diversity significance was calculated using a linear mixed-effects model (*lme4* and *lmerTest*) with sex, diet, and timepoint as main effects, diet:timepoint as an interaction, and subject as a random effect. This same model was used to test for significant changes in microbial species and functional pathway abundance with MaAsLin 3 at 5% FDR. Volcano plots, alpha diversity and Principal Coordinates Analysis (PCoA) plots were created with *ggplot2*. Taxa bar plots were created with the *fantaxtic* package.

### 2.6 Flow cytometry analysis

#### Evaluating circulating tumor burden

For routine monitoring of leukemia progression, whole blood was incubated with fluorochrome-labeled antibodies at 4°C for 20 minutes followed by RBC lysis with 1X RBC Lysis Buffer (Cat. #420301; BioLegend; San Diego, CA, USA) per manufacturer protocol prior to flow cytometry acquisition. Fluorochrome labeled-antibodies against CD19 (Clone 6D5, RRID AB_313642), CD45 (Clone 30-F11, RRID AB_2564590), and CD5 (Clone 53-7.3, RRID AB_312735) were obtained from BioLegend.

#### Flow cytometry acquisition and data analysis

Additional flow cytometry acquisition details are summarized in the **Supplementary File**. Flow cytometry gating strategies have been previously detailed (2).

### 2.7 Statistical analysis

Statistical analyses were performed using GraphPad Prism v 10.6.1 (GraphPad Software; Boston, MA, USA) and SAS software version 9.4 (SAS Institute Inc.; Cary, NC, USA). Data are reported as mean ± standard error of the mean (SEM) unless otherwise indicated. Statistically significant differences between two groups were determined using an unpaired Mann-Whitney U tests. Kaplan-Meier survival curves were generated using SAS PROC LIFETEST and univariable and multivariable Cox proportional hazard analysis using SAS PROC PHREG. *P* values less than 0.05 were considered statistically significant.

## 3 Results

### 3.1 Disease burden in peripheral blood and spleen

To characterize diet-dependent modifications to the gut microbiome during CLL initiation and progression, we utilized a diet modulation model in tandem with the Eµ-TCL1 adoptive transfer model of CLL. The adoptive transfer model reliably establishes CLL disease within recipient mice and closely recapitulates an aggressive form of CLL with rapid onset culminating in fatal disease within weeks, providing a way to characterize disease progression in respect to diet modification (**Fig. 1**). Following diet shift and CLL engraftment, mice were monitored until becoming moribund from their disease. CLL disease burden within the peripheral blood was not significantly different between the HF and CH mice across the monitored timepoints (**Fig. 2A**). We next explored the immune microenvironment within the spleen at end of study, given its role in forming an immunosuppressive tumor niche and major compartment for CLL development (28). F-HF mice displayed significantly greater disease burden in the spleen compared to F-CH mice at the time of sacrifice (**Fig. 2B**). There was a similar increase in disease burden within the spleen of M-HF mice compared to M-CH, though this was not statistically significant (**Fig. 2B**). Pooled analysis of all mice demonstrated significantly greater disease burden in spleen of mice on a HF diet compared to those on a CH diet (**Fig. 2B**).

**Figure 1.**
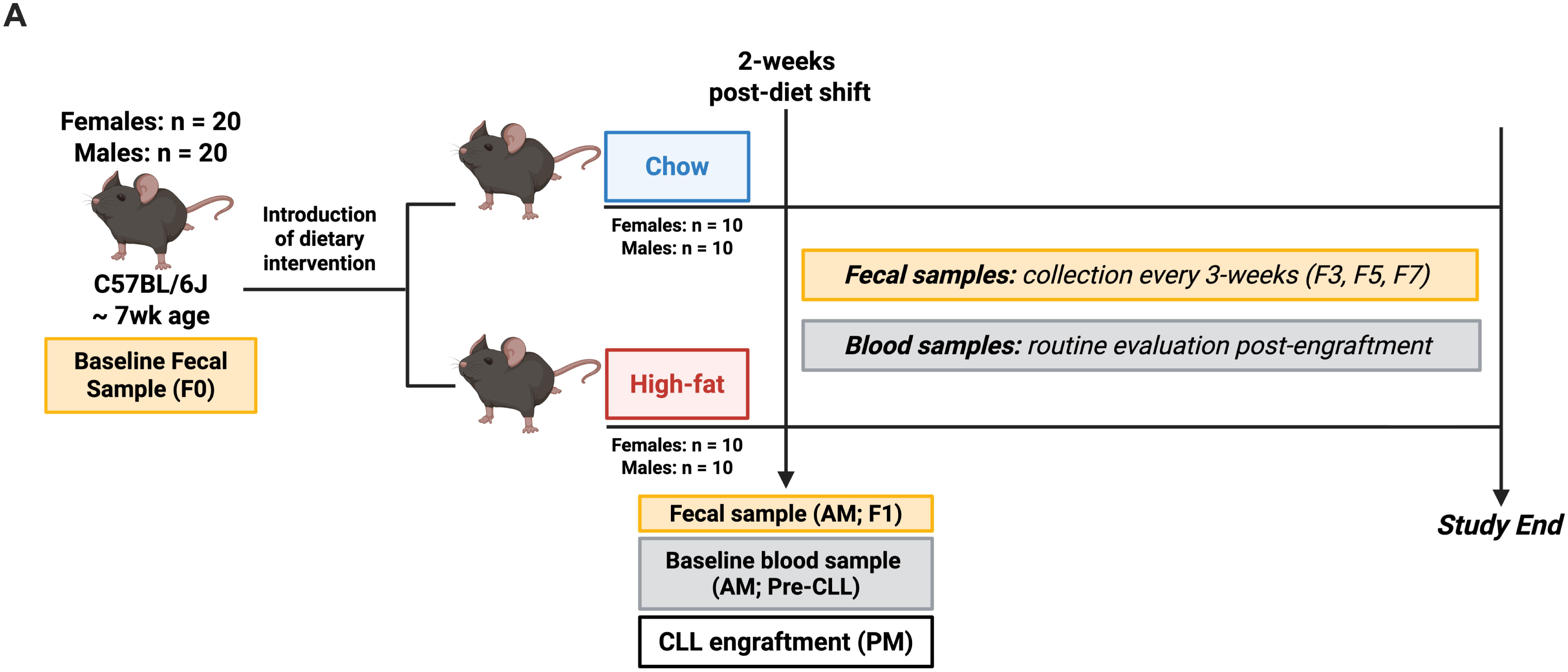
Diet modulation study outline. **A)** Study schematic illustrating the diet modulation model used. Prior to any study intervention, a baseline fecal sample (F0) was collected from all study mice (female, n = 20; male, n = 20). Subsequently, two-weeks prior CLL engraftment, wild-type (C57BL/6J) female and male mice were randomized into diet groups: female chow (F-CH; n = 10), female high-fat (F-HF; n = 10), male chow (M-CH; n = 10), and male high-fat (M-HF; n = 10). Weight of mice and food intake was monitored every ten days; water consumption was also monitored every ten days. Two-weeks post-diet shift, but prior CLL engraftment, a fecal sample (F1) and baseline (pre-CLL) blood samples were collected from all study mice. Briefly, all diet groups (n = 40) were engrafted via tail vein injection with 0.5e^7^ Eµ-TCL1 spleen-derived lymphocytes from a moribund Eµ-TCL1 mouse (≥ 90% CD45+/CD19+/CD5+ peripheral blood and spleen lymphocytes). Subsequent fecal samples were collected every three weeks (F3, F5, F7). Leukemic disease burden was routinely evaluated via flow cytometry. Due to overall survival of female and male mice, female flow analyses are documented at pre-CLL, 2-weeks post-engraftment, and 4-weeks post-engraftment. Male flow analyses are documented at pre-CLL, 2-weeks post-engraftment, 4-weeks post-engraftment, 8-weeks post-engraftment, and 12-weeks post-engraftment. Mice were followed until they became moribund, as outlined in the methods, and were subsequently sacrificed.

**Figure 2.**
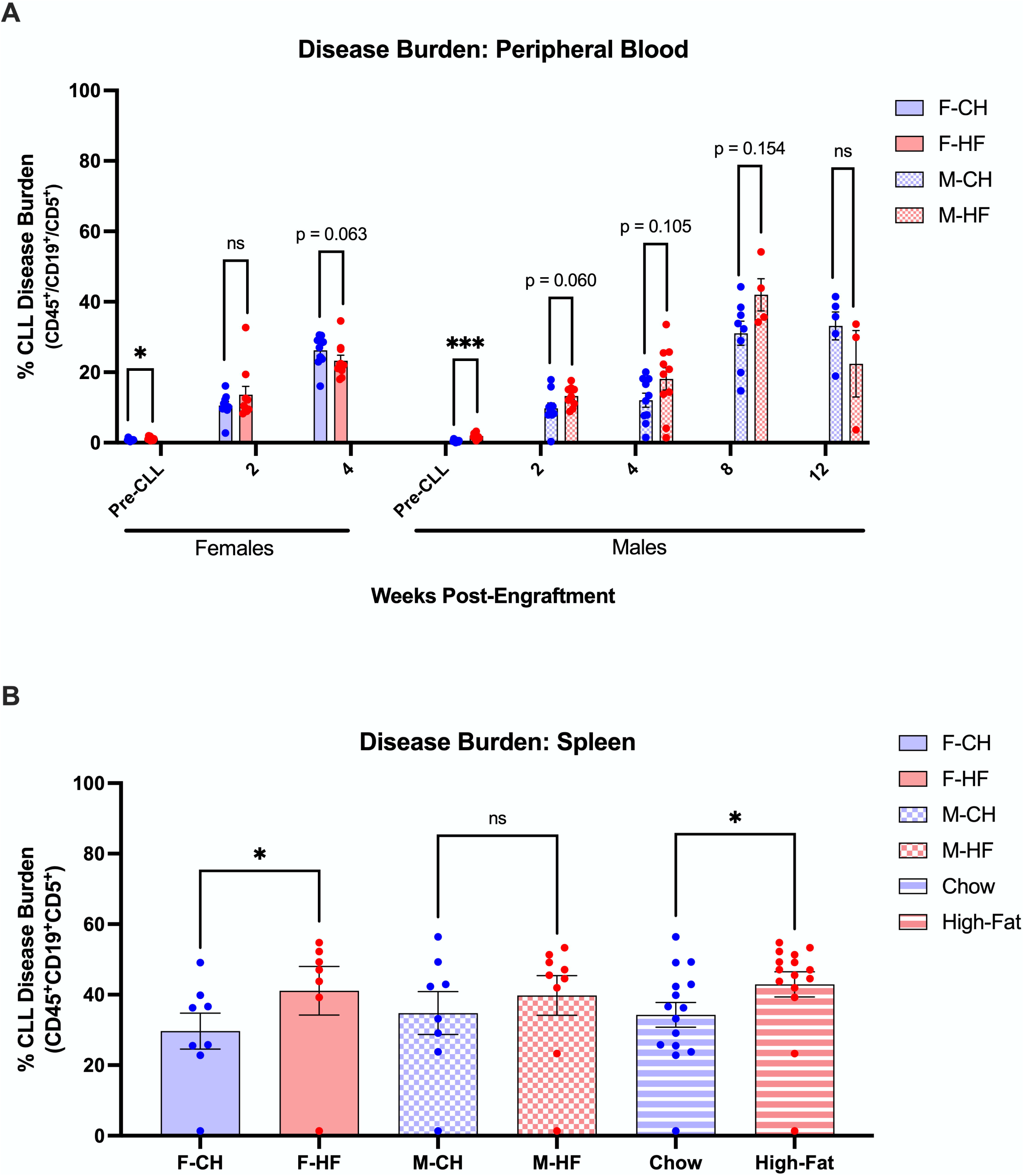
Expansion of CLL in the peripheral blood and spleens of diet study mice. **(A)** CLL disease burden was measured by percentage of CD45+/CD19+/CD5+ cells in the peripheral blood via flow cytometric analysis beginning prior CLL engraftment. Females were additionally monitored at 2-weeks post-engraftment and 4-weeks post-engraftment (F-CH, n = 10; F-HF, n = 10). Males were additionally monitored at 2-weeks post-engraftment, 4-weeks post-engraftment, 8-weeks post-engraftment, and 12-weeks post-engraftment (M-CH, n = 10; M-HF, n = 10). Asterisks denote the significance between diets at each time point (* *p* < 0.05, *** *p* < 0.001). Unpaired, Mann-Whitney U tests were used to determine significance between two groups. **(B)** CLL disease burden was measured by percentage of CD45+/CD19+/CD5+ cells in the spleen via flow cytometric analysis at end of study (i.e., when mice became moribund and sacrificed). Asterisks denote the significance between diets at each time point (* *p* < 0.05). Unpaired, Mann-Whitney U tests were used to determine significance between two groups. F-CH, female chow; F-HF, female high-fat; M-CH, male chow; M-HF, male high-fat.

### 3.2 T-cell populations in peripheral blood and spleen

All mice, regardless of diet assignment, showed a decrease in overall percentage of T-cells in the peripheral blood as CLL disease progressed (**Supplementary Figure S3A**). Further characterization of T-cells into CD4+ and CD8+ T-cells showed a non-significantly greater percentage of CD4+ T-cells in F-HF mice compared to F-CH over the course of the study. M-HF mice displayed a modestly greater percentage of CD4+ T-cells compared to M-CH mice at early study time points (i.e., 2-weeks and 4-weeks post-engraftment; **Supplementary Figure S3B**). M-HF mice featured modestly reduced percentages of CD8+ T-cells in the peripheral blood at early timepoints compared to M-CH mice that resolved over time. (**Supplementary Figure S3C**). F-HF mice showed a non-significantly greater percentage of CD8+ T-cells compared to F-CH mice at the 4-weeks post-engraftment timepoint (**Supplementary Figure S3C**).

T-cell dysfunction, marked by persistent co-expression of inhibitory receptors and impaired anti-tumor function, is a hallmark characteristic of CLL (28,29). No significant differences were observed in the co-expression of inhibitory receptors PD-1, TIM-3, and LAG3 on CD4+ or CD8+ T-cells within the peripheral blood of diet study mice (**Supplementary Figure S4A, B**). Lastly, to further characterize the extent of dysfunction, proportions of progenitor exhausted T-cells (T_PEX_; PD-1^int^/TIM-3^lo/-^) and terminally exhausted T-cells (T_TEX_; PD-1^hi^/TIM-3^hi^) were assessed (30,31).

Minimal significant differences were found in either population of CD4+ and CD8+ cells for both female and male diet study mice within the peripheral blood (**Supplementary Figure S5A-D**).

Despite no overt changes observed in the peripheral blood, we looked to characterize T-cell subsets in within the spleen. Similarly, minimal differences in T-cell populations and exhaustion phenotypes were observed within the splenic microenvironment (**Supplementary Figure 6-8)**. Overall, we note that dietary assignment did not significantly influence the circulating or splenic T-cell subsets in Eµ-TCL1 adoptive transfer mice.

### 3.3 Impact of diet on survival outcomes in adoptive transfer Eµ-TCL1 mice

We next investigated the impact of diet modulation on overall survival. An overall reduction in survival was observed for the HF-diet mice compared to CH-diets (**Fig. 3**, *p* <0.001). F-HF mice displayed significantly reduced overall survival compared to F-CH mice. Specifically, F-CH mice had a median survival of 46 days compared to F-HF mice with a median survival of 31 days (**Fig. 3**). Similarly, M-HF mice lived significantly fewer days than M-CH mice (**Fig. 3**). Of note, male mice lived significantly longer than female mice in this study (**Fig. 3**), consistent with published literature regarding the Eµ-TCL1 model of CLL (32,33). On multivariable analysis using a Cox proportional hazard model (CPH), both female sex (hazard ratio (HR) 0.19, *p* < 0.001) and HF diet (HR 0.30, *p* = 0.002) were negatively associated with survival (**Supplementary Table 2**). Additional analysis using the CPH model identified a potential interaction between sex and diet, where the impact of the HF diet was worse on female mice vs. males (*p* = 0.0020) as presented in **Supplementary Table 2**. In addition, scatter plots of days on study by gender and diet type revealed differing slopes in log-transformed survival times between CH and HF diet (**Supplementary Fig. S2**). Notably, female mice on the HF diet demonstrated a steeper decline, indicating accelerated mortality impact from the HF-diet compared with male mice.

**Figure 3.**
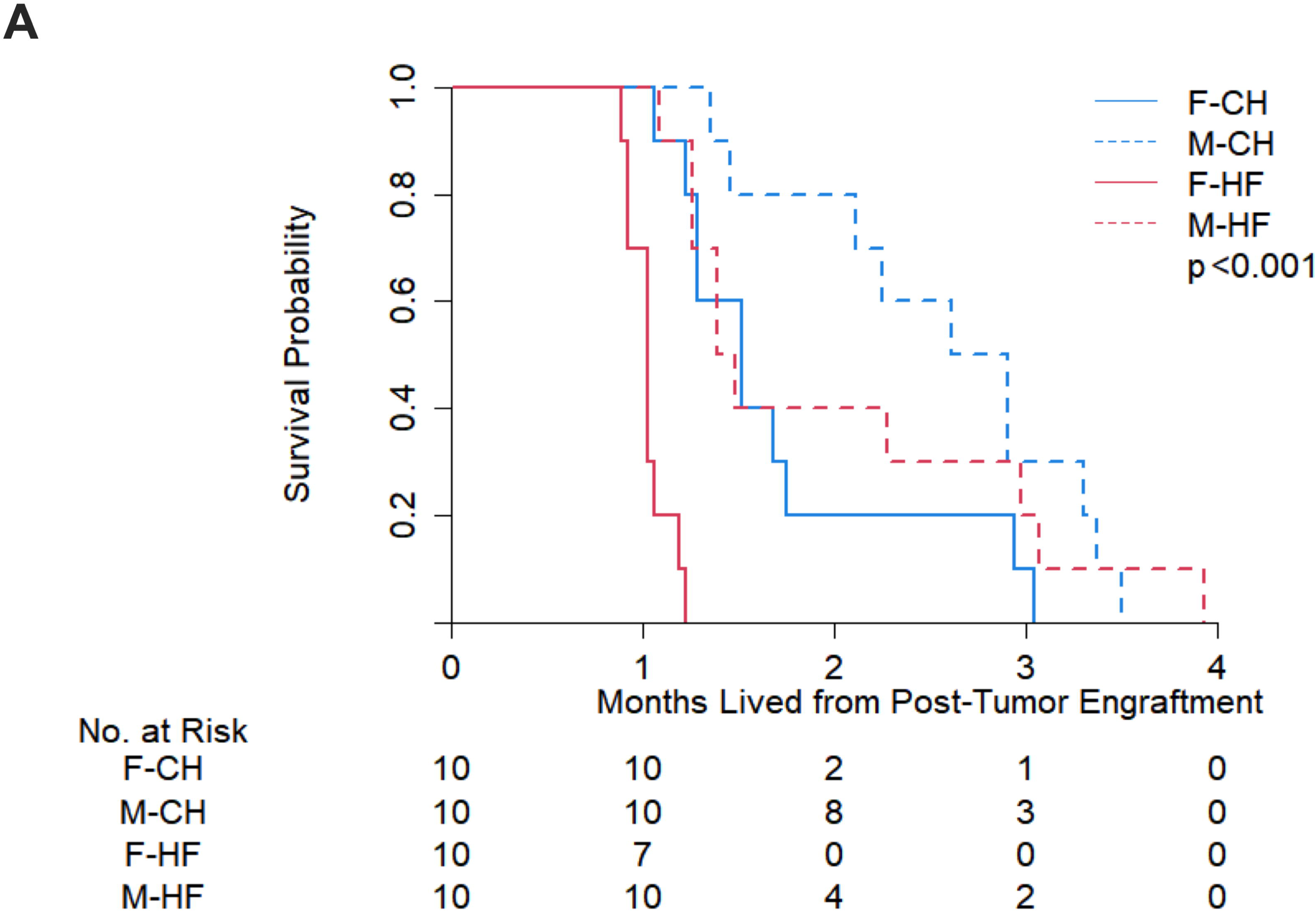
Effect of diet modulation on CLL survival. **(A)** Kaplan-Meier survival analysis of all diets over the course of the study (F-CH, n = 10; F-HF, n = 10; M-CH, n = 10; M-HF, n = 10). *P*<0.001. Calculated median survival time (days) is provided in the figure legend. Multivariable Cox proportional hazards model comparing the effects of gender and diet on days-to-event is provided in **Supplementary Table 2**. F-CH, female chow; F-HF, female high-fat; M-CH, male chow; M-HF, male high-fat.

### 3.4 Microbiome analysis of diet study mice on chow and high-fat diets

#### Microbial Population Changes

We next examined the impact of each diet on the gut microbiome. Exposure to the HF diet was associated with an immediate reduction in alpha diversity compared to CH diet, a difference that was sustained throughout the study (**Fig. 4A**). Beta-diversity, measured using Bray-Curtis dissimilarity notes that although the samples were similar at baseline between the HF and CH diets, an immediate change was similarly identified and sustained in beta-diversity (**Fig. 4B and 4C**), indicating significant shifts in gut microbial composition between diets.

**Figure 4.**
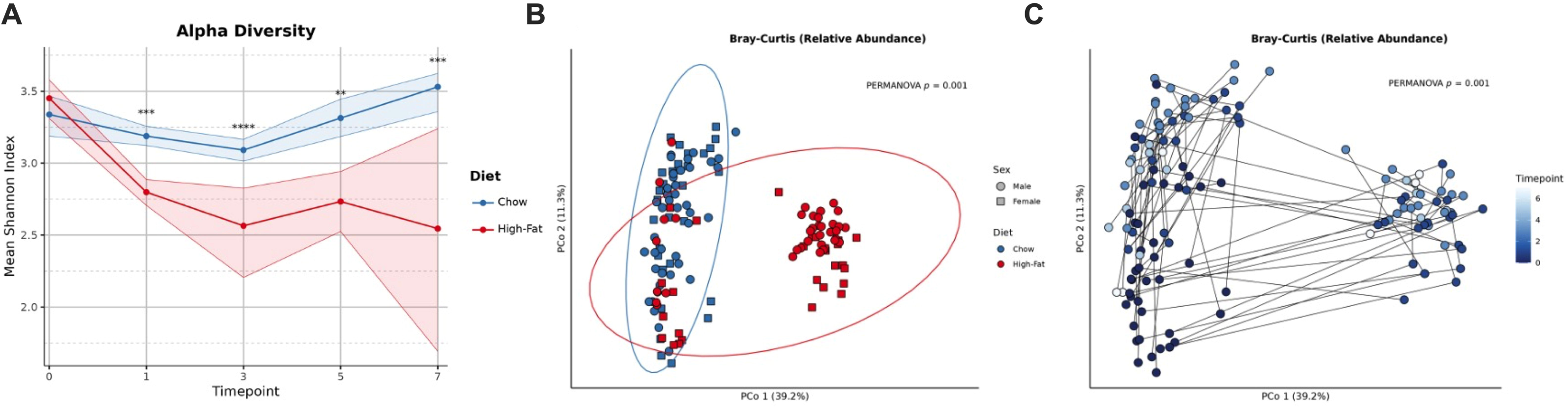
Effect of diet modulation on microbiome alpha and beta diversity. **(A)** Alpha diversity measured by Mean Shannon Index between CH and HF diets. Alpha diversity measures are not separated by sex. Data are represented as a line chart. Asterisks denote the significant diet:timepoint interactions in the HF group with respect to CH at each time point (* *p* < 0.05, ** *p* < 0.01, *** *p* < 0.001). A linear mixed-effects model was applied for significance testing. **(B)** PCoA plot depicting Bray-Curtis dissimilarity as a measure of beta diversity between M-CH, M-HF, F-CH, and F-HF. Male mice are denoted with either blue circles (M-CH) or red circles (M-HF). Female mice are denoted with either blue squares (F-CH) or red squares (F-HF). PERMANOVA was applied to test for differences in beta diversity between subjects. **(C)** PCoA plot depicting Bray-Curtis dissimilarity as a measure of beta diversity between study timepoints. Male timepoints include pre-diet shift, pre-CLL engraftment, 4-weeks post-engraftment, 8-weeks post-engraftment, and 12-weeks post-engraftment. Female timepoints include pre-diet shift, pre-CLL engraftment, and 4-weeks post-engraftment. Data points are not specified by diet or sex. PERMANOVA was applied to test for differences in beta diversity within subjects.

We next characterized which specific microbiota change by examining both microbiome population changes (**Fig. 5**) and metabolic pathway changes (**Fig. 6**).The relative abundance taxa bar plot (**Fig. 5**) demonstrates that HF diet consumption led to a sustained increase in the presence of *Akkermansia muciniphila*, mixed changes in unclassified *Bacteroidota* (synonym *Bacteroidetes*) phylum members, a reduction in *Firmicutes* (other) members, and a loss of *Tenericutes* species *SGB44562* (**Fig. 5A**). We performed a differential abundance analysis to better determine the specific species affected by dietary assignment, sex, and time in study mice. The heatmap provided (**Fig. 5B**) confirms that the HF diet across timepoints led to statistically significant increases in *A. muciniphila*, *Bacteroidetes thetaiotaomicron*, and *Lachnospiraceae_GGB28916,* with reductions in *Acutalibacter muris* and multiple *Lachnospiracaeae*, *Oscillospiraceae*, and Muribaculaceae family members (**Fig. 5B**). Separately, the impact of diet (**Fig. 5C**) and sex (**Fig. 5D**) on differentially abundant bacterial taxa is displayed by volcano plots noting the magnitude in fold-abundance changes in the taxa differences observed in the heatmap. Female sex (**Fig. 5D**) was associated with increases in *Lachnospiraceae* family and *Firmicutes* phyla members with reductions in *Muribaculaceae* and *Pumilibacteraceae* family members (**Fig. 5D**). Additional taxa differentially abundant in the volcano plots are depicted in the **Supplementary Data File**.

**Figure 5.**
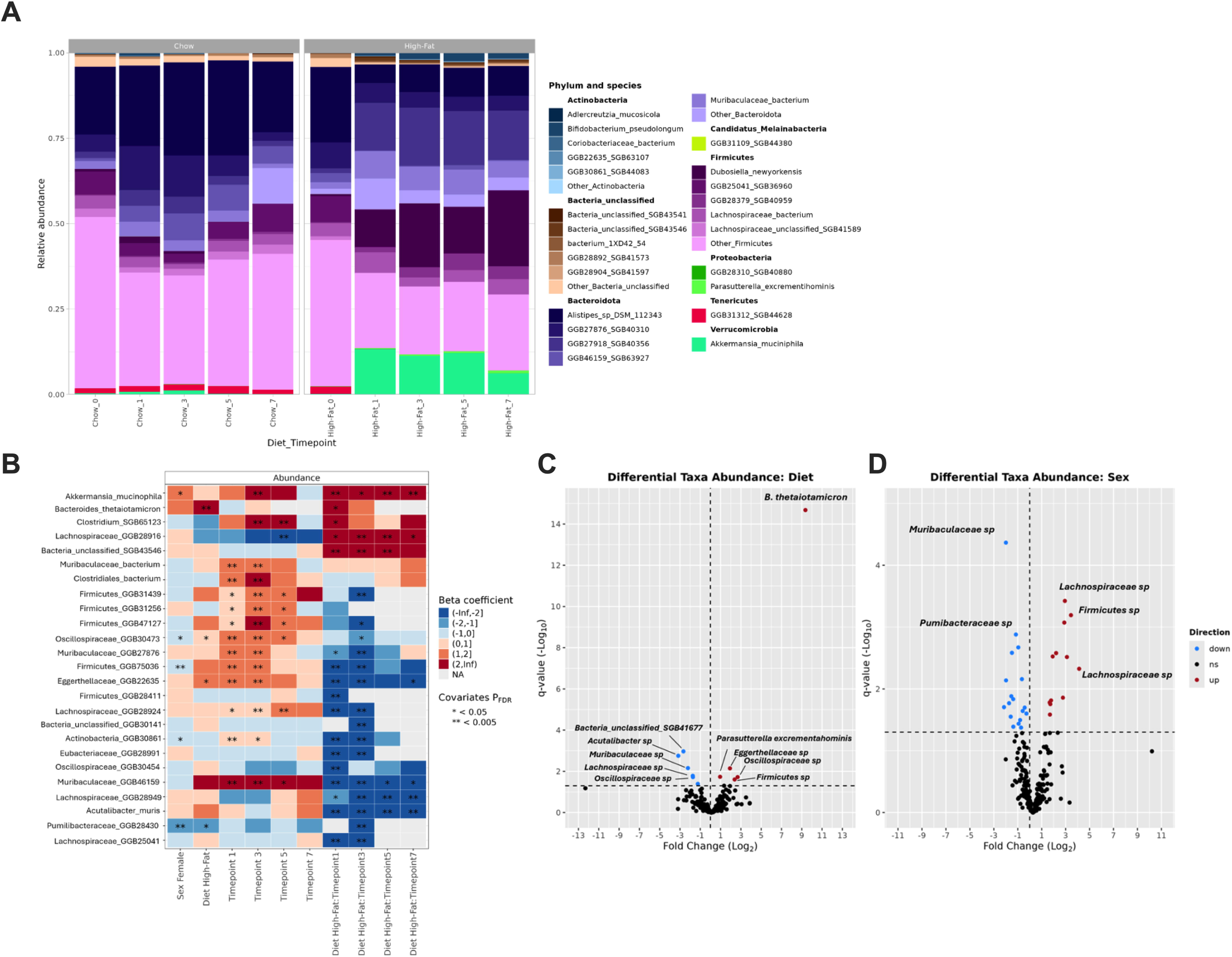
Microbial species that differ significantly in abundance between diet groups and sexes. **(A)** Relative abundance bar plots of bacterial taxa present at the species level in CH and HF diets at pre-diet shift, pre-CLL engraftment, 4-weeks post-engraftment, 8-weeks post-engraftment, and 12-weeks post-engraftment. The five most abundant species within each phylum are displayed, with remaining species aggregated into an “Other” category. **(B)** Heatmap depicting differential taxa abundance at the genus-species level. Labels along the x-axis correspond to regression coefficients (i.e., independent variables) in a fitted MaAsLin 3 model, with colors denoting the sign and magniture of those coefficients. **(C)** Volcano plot depicting downregulated (blue dots) and upregulated (red dots) taxa based on diet (i.e., CH vs. HF). **(D)** Volcano plot depicting downregulated (blue dots) and upregulated (red dots) taxa based on sex (i.e., male vs. female). CH, chow; HF, high-fat; pre-diet shift = Chow/High-Fat_0; pre-CLL engraftment = Chow/High-Fat_1; 4-weeks post-engraftment = Chow/High-Fat_3; 8-weeks post-engraftment = Chow/High-Fat_5; 12-weeks post-engraftment = Chow/High-Fat_7. Volcano plot p-values and fold changes were outputted by MaAsLin 3.

**Figure 6.**
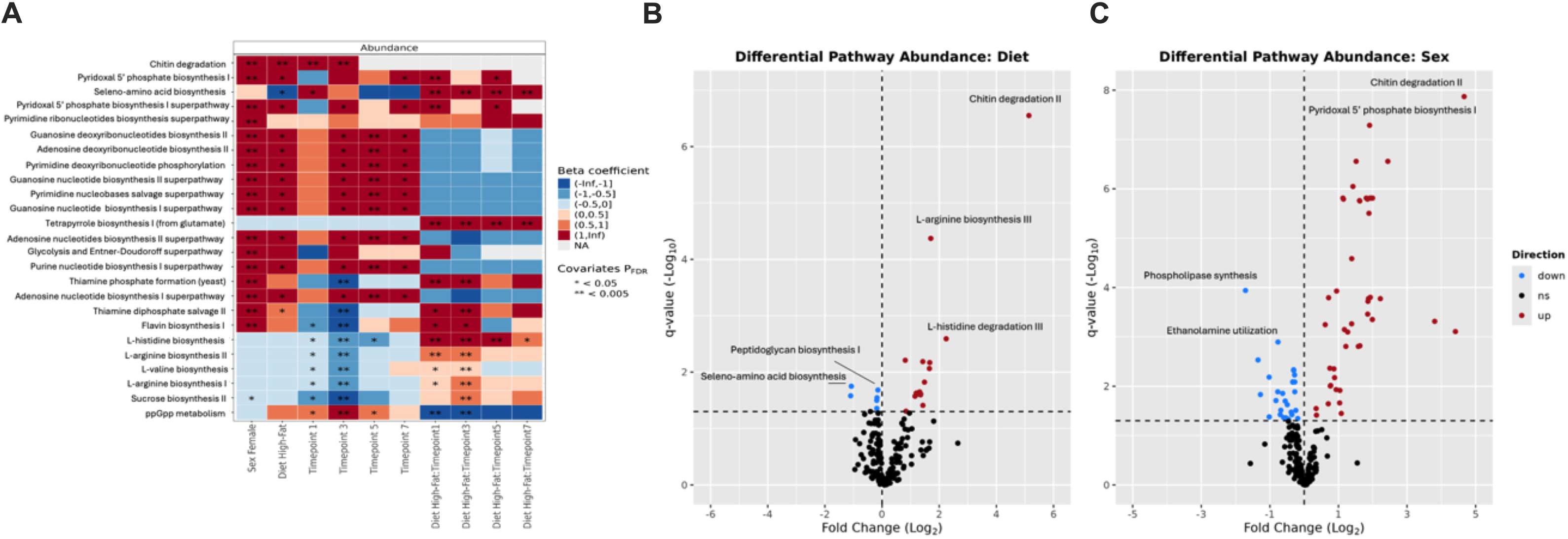
Microbiome functional pathways that differ significantly in abundance between diet groups and sexes. **(A)** Heatmap depicting differential abundanct MetaCyc pathways. Labels along the x-axis correspond to regression coefficients (i.e., independent variables) in a fitted MaAsLin 3 model, with colors denoting the sign and magniture of those coefficients. **(B)** Volcano plot depicting downregulated (blue dots) and upregulated (red dots) pathways based on diet (i.e., CH vs. HF). **(C)** Volcano plot depicting downregulated (blue dots) and upregulated (red dots) pathways based on sex (i.e., male vs. female). Volcano plot p-values and fold changes were outputted by MaAsLin 3.

#### Functional Metabolic Pathway Changes

Following the microbial classification and abundance measurements, we next evaluated the impact of diet on the abundance of microbiome functional pathways using the metagenomic data obtained from DNA sequencing (**Fig. 6**). When compared to CH diet compiled across all timepoints, the HF diets was associated with increases in genes for chitin degradation and multiple nucleotide biosynthesis pathways including pyrimidine bases, guanosine, and adenosine (**Fig. 6A)**. A reduction in seleno-amino acid synthesis genes was also observed. In a comparison to CH diets timepoint-by-timepoint, increases across all timepoints were observed for genes related to tetrapyrrole synthesis (from glutamate) and histidine biosynthesis. Early timepoints 1 and 3 demonstrated that HF diets were associated with increases in multiple arginine biosynthesis pathways, valine biosynthesis, and a reduction in ppGpp synthesis (**Fig. 6A**). Volcano plots highlight the fold changes in each pathway globally for the HF diet compared to CH at timepoint 0 as control (**Fig. 6B**). When analyzing by sex (**Fig. 6C**), multiple pathways were enriched including chitin degradation and multiple nucleotide biosynthetic pathways (**Fig. 6C**). Reductions in genes for phospholipase synthesis and ethanolamine utilization were also noted. A complete table of enriched and reduced metabolic pathways demonstrated on the volcano plots is provided in the **Supplementary Data File**.

## 4 Discussion

To our knowledge, the data presented herein provides one of the first comprehensive analyses of the direct effect of diet modulation on the gut microbiome, disease progression, and survival in the pathogenesis of CLL, using an established murine model. HF diet was associated with profound dysbiotic changes to the gut microbiome with a reduction in community alpha diversity and significant differences in beta diversity compared to control CH diet. These observations highlight the profound, immediate, and sustained impact a diet enriched in fat and carbohydrates, and low in fiber (designed to mimic the “Western diet”) had on the gut microbiome within this model. Although multiple shifts in taxa were observed in this analysis, several observations stand out. First is the direct effect of a HF diet on alpha diversity. Low alpha diversity has been negatively associated with both the pathogenesis and outcomes across many oncologic diseases including CLL (34–36). For patients receiving cellular therapy or immunotherapy, reductions in alpha diversity within the gut microbiome have been consistently associated with inferior survival outcomes – highlighting that perturbations to the gut microbiome may cause consistent and robust impacts to oncologic disease (37–40). The majority of studies demonstrating this relationship between low diversity and poor clinical outcomes are typically limited in their ability to identify key drivers of this loss in diversity aside from the disease under study or antibiotic exposure. Our study adds to the growing body of literature linking gut microbiome diversity to clinical outcomes in oncologic disease generally and CLL specifically by demonstrating how a high fat, high-carbohydrate diet can significantly influence the diversity of the gut microbiome.

Second, we observed significant population and metabolic pathway changes related to consumption of HF diet. The strongest and most consistent signal was the sustained increase in *A. muciniphila* and *B. thetaiotaomicron*. Notably, *A. muciniphila* has been characterized in the gut microbiome in multiple studies, yet its role as a beneficial or pathogenic commensal is uncertain (41). Functionally, it is known to be a key mucin degrader, which may weaken the gut lumen barrier and provide an opportunity for pathogenic or inflammatory species to activate host immune responses via direct translocation or cytokine production (42–44). Alternatively, *A. muciniphila* has also been associated with favorable responses to immunotherapy (45). Ultimately, although our study cannot identify a mechanism linking *A. muciniphila* to CLL pathogenesis, its sustained increase in the context of a diet intervention study in CLL provides a compelling foundation for further analysis, particularly investigating its role as a mucin degrader at the gut lumen-blood barrier. *B. thetaiotaomicron* was also observed to be significantly increased in the microbiome of CLL mice receiving the HF diet. While a known commensal organism in the gut microbiome, the beneficial or harmful impacts of *B. thetaiotaomicron* are also mixed in the literature, supporting further evaluation of its role in CLL pathogenesis (46).

Third, we observed that the community shifts observed in microbiome populations led to changes in the metabolic functional potential of the gut microbiome. HF diets resulted in enrichment of diverse metabolic pathways related to arginine and histidine biosynthesis, chitin degradation, and nucleotide biosynthesis (47). Although the functional consequences of these enriched metabolic pathways are unclear in this analysis, several potential hypotheses could explain their significance.

Notably, arginine and amino acid metabolism plays an important role in the CLL tumor microenvironment, where prior work has identified the need for arginine and other amino acids to promote CLL pathogenesis (48–50). One potential explanation for our observations that link HF diet to the gut microbiome and CLL pathogenesis is that changes in the community profile (i.e., *A. muciniphila* growth) may alter the GI lumen barrier, allowing enhanced metabolic exchange with sustaining metabolites provided by a dysbiotic gut microbiome. In addition, glutamate metabolism contributes to nitrogen recycling and has been previously identified as a means by which changes to the gut microbiome can influence the tumor microenvironment in multiple myeloma (51).

Aside from specific impacts of the HF diet alone, we also noted sex-specific differences in the gut microbiome and CLL survival. Importantly, the difference in survival in the female mice compared to male mice is a known characteristic of the Eµ-TCL1 CLL model, where male mice tend to develop disease later and on average live longer than female mice (32,33). Preclinical studies in the Eµ-TCL1 adoptive transfer model mice have shown female recipients succumbed to CLL earlier than male recipients (32,33). Although this likely occurs for reasons beyond the gut microbiome, we provide evidence for the first time to suggest that there are sex-specific differences in the gut microbiome occurring within this model, which require further exploration both in animal models and human subjects. Cox proportional hazards modeling identified a diet–sex interaction in survival outcomes, with female mice on the HF diet exhibiting worse outcomes than male mice. These findings support further large scale studies of sex specific microbiome differences in CLL. Additionally, HF diet significantly impacted overall survival in the Eµ-TCL1 CLL model compared to standard CH diet. Other studies have also demonstrated the negative impact of high-fat/Western diets on disease state in comparison to mice on a standard chow diet (52–54). The CH diet represents a natural ingredient diet that is used to fulfill the everyday needs of laboratory animal colonies. In essence, it’s a combination of unrefined whole grains, grain by-products, and fats at levels that are likely the most natural and healthiest diet for the mice. The alternative HF diet is notably different in both its macronutrient profile – high in fat and carbohydrates, and low in fiber – and manufactured as a processed diet. Although we cannot determine which nutrient (high fat, low fiber, or a combination) and/or manufacturing process caused the observed differences in microbiome and CLL disease burden, the effect of diet on CLL survival was striking. A study conducted by Solans *et al.* found a potential correlation between consumption of ultra-processed food and drinks (UPF) and CLL incidence (23).

Our study also suggests a potential mechanism between CLL pathogenesis and the gut microbiome. Although we did not observe a difference in CLL expansion within the peripheral blood, we noted greater CLL involvement within the spleen at time of sacrifice in mice receiving the HF diet. Consistent with the Eµ-TCL1 CLL model, mice developed progressive splenomegaly that significantly impairs their mobility, eventually prompting euthanasia. This was observed more rapidly in the mice consuming HF diet compared to those consuming CH diet, and surprisingly more evident in the female mice receiving the HF diet. Additional analysis is required to determine if the effect of diet exposure on CLL is driven via specific interaction within the spleen compartment as opposed to the peripheral blood, though others have observed that the gut microbiome’s impact on oncologic disease may occur via immune-related activity in the spleen (55).

Our study has several notable limitations. First, the limited number of animals utilized in the experiment for practical considerations may have limited our ability to statistically model the interaction between diet and sex variables, among other untested variables as well. As spleens were not prospectively collected during the experiment, we are unable to determine if CLL involvement in the spleen occurred more rapidly in the HF vs. CH diet. Although the cause of death/sacrifice was consistent with the known behavior of the Eµ-TCL1 mouse model due to progressive splenomegaly, we cannot rule out alternative causes of death related to the different diet exposures. The metabolic pathway analysis was based on functional potential provided by the genomic testing in this study and direct metabolites were not measured. In addition, dietary effects on other physiological or biological parameters could have affected survival through both CLL and non-CLL related activities that were not assessed in this experiment. Although the HF diet significantly impacted the gut microbiome and CLL survival, supporting our initial hypothesis, these relationships are associations at best, where additional experimentation, including in germ-free models, would be required to help prove causal links.

Ultimately, we conclude that exposure to the HF diet caused significant perturbations to the gut microbiome across community measurements of diversity, taxa abundance, and functional pathway abundance; and HF diet was associated with inferior survival in the Eµ-TCL1 murine model of CLL. These findings provide a firm foundation for further exploration regarding the role of diet in CLL pathogenesis and survival and suggest potential mechanisms to test, including worsening gut permeability and amino acid metabolism.

## Supporting information

Supplementary Date File

## 6 Author Contributions

S.A.S., J.B.H., C.R.D., and D.E. contributed to the conception, design, and development of methodology. S.A.S., A.L.S., and E.M.D. conducted the murine studies. S.A.S. and J.B.H. conducted the data analysis. J.B.H. analyzed the metagenomic shotgun sequencing data. S.A.S., J.B.H., J.B.C., A.K.S, S.N.R., C.R.D., and D.E. contributed to the analysis and interpretation of the data. S.A.S., C.R.D, and D.E. wrote the original manuscript draft and all authors contributed to the editing of the article and reviewed the manuscript. All authors provided final approval and agreed to be accountable for all aspects of the work, ensuring its integrity. C.R.D. and D.E. jointly managed and supervised all study aspects and obtained funding. All authors have read and agreed to the published version of the manuscript.

## 8 Acknowledgments

We acknowledge the UNMC Comparative Medicine Core for all animal housing and research technical support. We also thank Carlo M. Croce (The Ohio State University) for access to the Eµ-TCL1 mice. The authors are grateful to the UNMC Genomics Core and the UNMC Flow Cytometry Research Facility. The UNMC Genomics Core receives partial support from the NIGMS INBRE-5P20 GM103427-21 grant as well as the FPBCC Support Grant – 5P30 CA036727-36. The UNMC Flow Cytometry Research Facility is supported by state funds from the Nebraska Research Initiative and the FPBCC National Cancer Institute Cancer Support Grant (P30 CD036727).

## 10 Data Availability Statement

The sequencing data generated in this study are deposited in the Sequence Read Archive (SRA) at the National Center for Biotechnology Information (NCBI) using PRJNA##### and will be made public upon publication. The pipeline used to process shotgun sequence data is available at ^1^ (DOI: 10.5281/zenodo.17715736). The code used for the analysis of microbiome data is available at https://github.com/clayton-lab/cll_mouse_microbiome_manuscript. All other data generated in this study are available upon request from the corresponding authors.

