## Supplementary Materials for "Impact of high-fat Western diet on chronic lymphocytic leukemia disease progression and gut microbiome profile in Eµ-TCL1 mice"

### *Supplementary Material*

#### 1 Supplementary Data

##### 1.1 Flow cytometry analysis

*Immunophenotyping of peripheral blood lymphocytes:* At study end, a comprehensive assessment of T-cells was conducted. Briefly, ~25 µL whole blood was incubated with fluorochrome-labeled antibodies at 4°C for 20 minutes followed by red blood cell lysis with 1X RBC Lysis Buffer (Cat. #420301; BioLegend; San Diego, CA, USA) per manufacturer protocol prior to flow cytometry acquisition. Fluorochrome-labeled antibodies against CD19 (Clone 6D5, RRID AB_313642), CD4 (Clone RM4-5, RRID AB_893326), CD45 (Clone 30-F11, RRID AB_2564590), CD8 (Clone 53-6.7, RRID AB_2563057), PD-1 (Clone 29F.1A12, RRID AB_2562568), and LAG3 (Clone C9B7W, RRID AB_10639727) were obtained from BioLegend. The fluorochrome-labeled antibodies against CD3 (Clone 145-2C11, RRID AB_2870231) and TIM-3 (Clone 5D12, RRID AB_2744186) were obtained from BD Biosciences.

*Immunophenotyping of splenic immune populations:* The protocol detailed below was utilized to label extracellular markers for flow cytometric analysis of splenocytes. *Live/Dead staining:* Approximately $3e^6$ cells were suspended in 100 µL FACS buffer (PBS + 2% hi-FBS) containing 0.3 µL Live/Dead Near IR fixable dye (BioLegend). Cells were vortexed and incubated at 4°C for 20 minutes, protected from light. Following incubation, cells were washed with 1 mL PBS, centrifuged at 350 x g for 5 minutes, and supernatant was decanted. *Fc Receptor blocking:* Following Live/Dead staining, cells were resuspended in 50 µL FACs buffer containing 2 µL TruStain FcX™ (BioLegend), vortexed, and incubated at 4°C for 5-10 minutes, protected from light. *Extracellular staining:* Immediately following incubation, and without washing 100 µL antibody mix in FACS buffer was added to each sample. Samples were vortexed and incubated at 4°C for 30 minutes, protected from light. Following incubation cells were washed with 1 mL PBS, centrifuged at 350 x g for 5 minutes, and supernatant was decanted. Fluorochrome-labeled antibodies against CD19 (Clone 6D5, RRID AB_313642), CD4 (Clone RM4-5, RRID AB_893326), CD45 (Clone 30-F11, RRID AB_2564590), CD5 (Clone 53-7.3, RRID AB_312735), CD8 (Clone 53-6.7, RRID AB_2563057), PD-1 (Clone 29F.1A12, RRID AB_2562568), and LAG3 (Clone C9B7W, RRID AB_10639727) were obtained from BioLegend. The fluorochrome-labeled antibodies against CD3 (Clone 145-2C11, RRID AB_2870231) and TIM-3 (Clone 5D12, RRID AB_2744186) were obtained from BD Biosciences. *Fixation:* Cells were resuspended in 300 µL Fixation Buffer (BioLegend), vortexed, and incubated for 30 minutes at room temperature. Following incubation, cells were washed twice with 2 mL PBS and resuspended in 300 µL FACS Buffer. Samples were stored at 4°C, protected from light, until flow cytometric data acquisition.

*Flow cytometry acquisition and data analysis:* Flow cytometry was performed on a LSRFortessa X-50 (BD Biosciences; San Jose, CA, USA). Single lymphocytes were gated by forward and side scatter. Live cells were gated by Live/Dead Near IR (Invitrogen; Waltham, MA, USA) negative staining. Correct compensation, proper acquisition setup, and positive gates based on appropriate fluorescence minus one (FMO) controls were ensured. Data were analyzed using Kaluza v2.1 (Beckman Coulter; Brea, CA, USA). Results are expressed as the proportion of cells expressing antigens of interest (percent gated positive).

#### 2 Supplementary Figures and Tables

##### 2.1 Supplementary Figures

**Supplementary Figure S1**

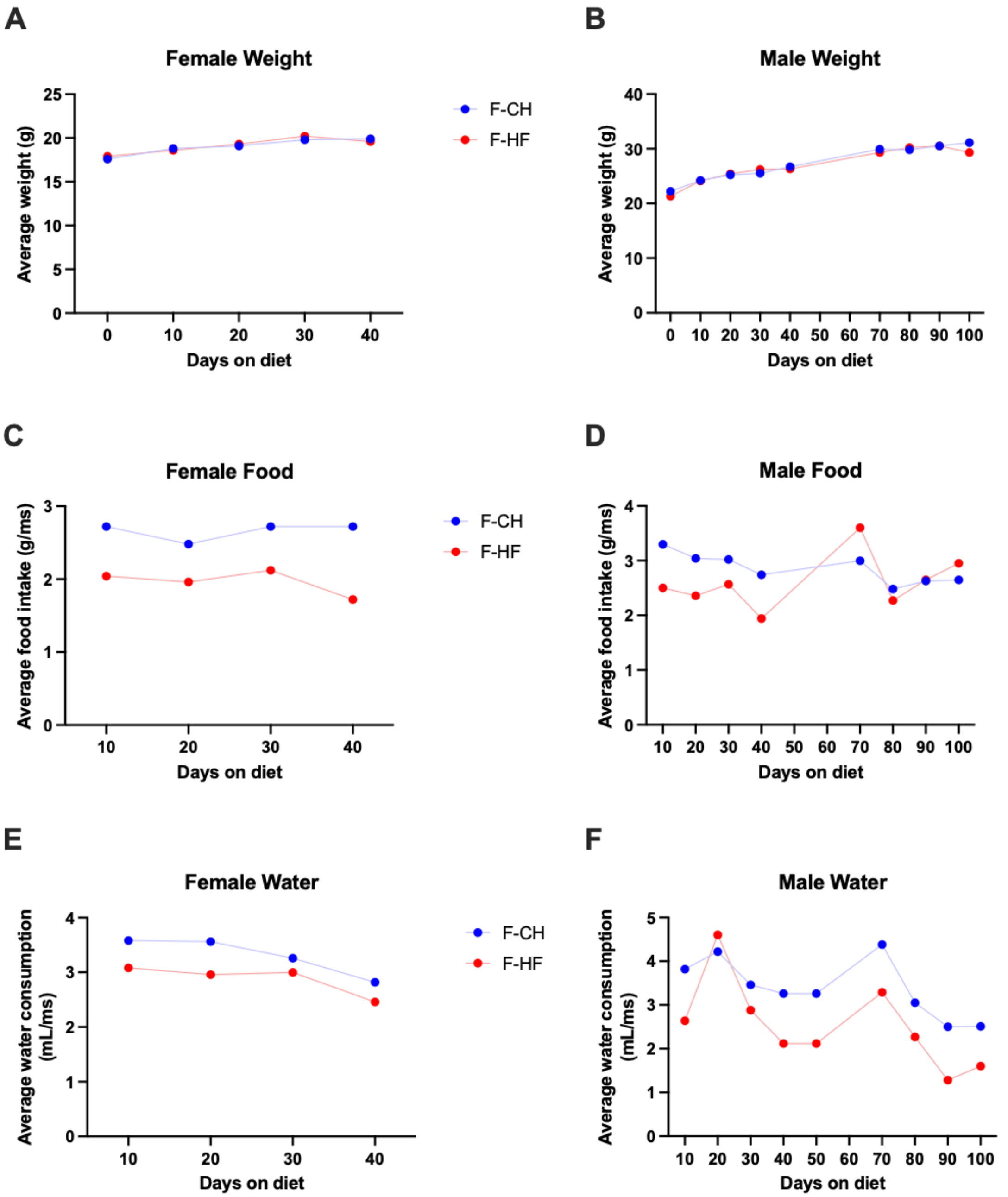


**Supplementary Figure S1. Confirmation of diet and water intake in study mice. (A, B)** Average weight (g) of mice enrolled per diet for females **(A)** and males **(B)**. For each diet, n = 10 mice were separated into two cages. Average weight values for each diet were calculated by weighing all ten mice between the two cages and averaging the values. The weight of mice was monitored every ten days.

**(C, D)** Average food intake (g) of mice enrolled per diet for females **(C)** and males **(D)**. Briefly, at the start of the diet intervention, the weight of food given per cage was documented. Every ten days the remaining food (loose and confined) in the cage was weighed and documented. The amount of food consumed in each cage was averaged, considering the number of days passed and number of mice remaining in the cage. Final average values were graphed. Inconsistencies in amount of food consumed varies across diets based on the amount of food replaced each time and the number of mice per cage (i.e., as mice were sacrificed or succumbed to disease). **(E, F)** Average water consumption (mL) of mice enrolled per diet for females **(E)** and males **(F)**. Briefly, at the start of the diet intervention, the volume of water given per cage was documented. Every ten days, the remaining water in the cage was measured and documented. The amount of water drank in each cage was averaged, considering the number of days passed and number of mice remaining in the cage. Final average values were graphed. Inconsistencies in the amount of water drank varies across diets based on the number of mice per cage (i.e., as mice were sacrificed or succumbed to disease).

**Supplementary Figure S2**

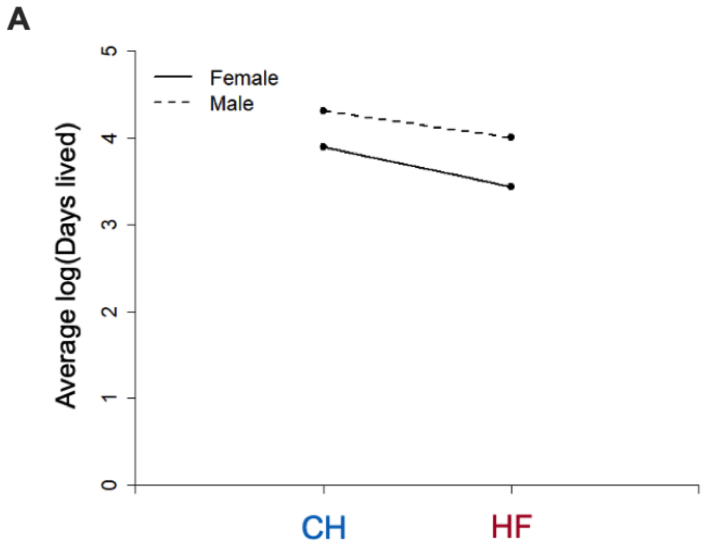


**Supplementary Figure S2. Scatter plot of days on study stratified by gender and diet.** Slopes of the log-transformed survival times (days lived) differ between diet groups (CH vs. HF) and sex, demonstrating that female mice on a HF diet experienced accelerated mortality compared to male mice on a HF diet.

#### Supplementary Figure S3

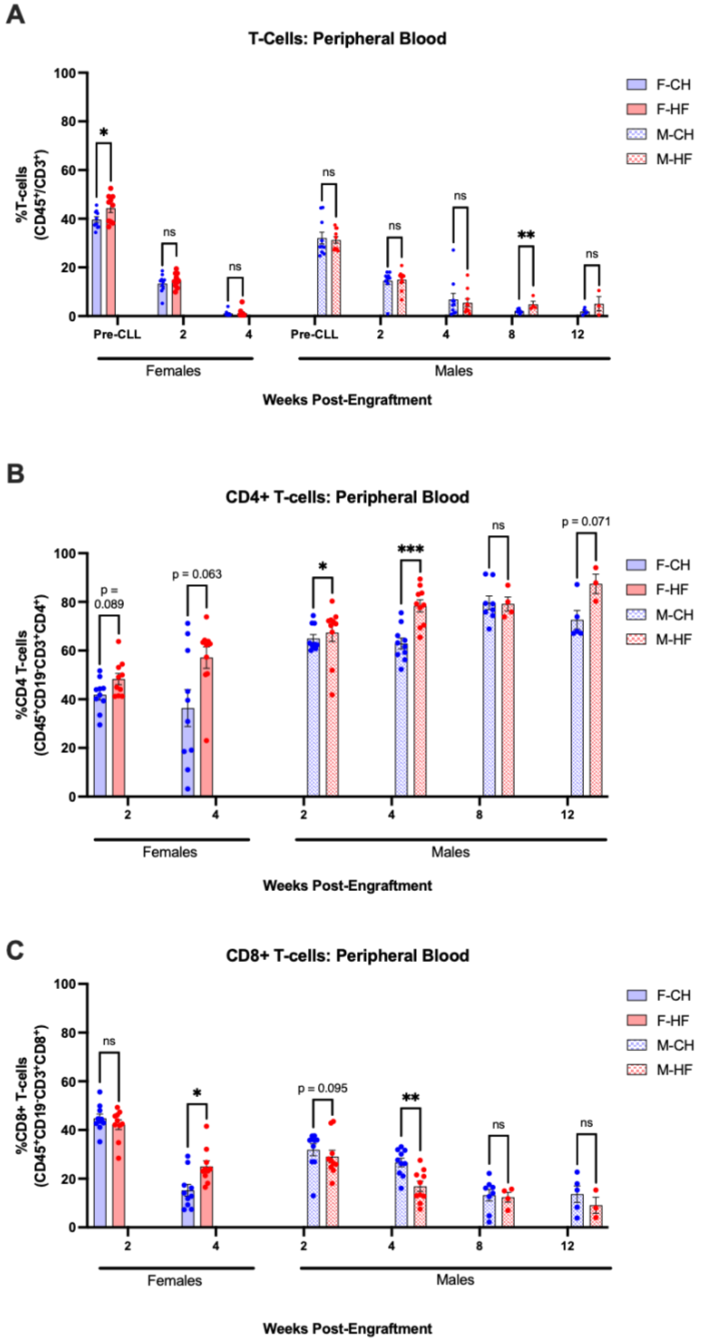


**Supplementary Figure S3. Characterization of T-cell subsets in the peripheral blood of female and male diet study mice. (A)** Percentage of T-cells found in the peripheral blood of diet study mice (F-CH, n = 10; F-HF, n = 10; M-CH, n = 10; M-HF, n = 10). T-cells were gated as CD45+/CD19-/CD3+. Asterisks denote the significance between either F-CH and F-HF mice or M-CH and M-HF mice (* $p < 0.05$, ** $p < 0.01$, ns = not significant). Unpaired Mann-Whitney U t-test was applied for testing. **(B)** Percentage of CD4+ T-cells found in the peripheral blood of F-CH, F-HF, M-CH, and M-HF diet study mice. CD4+ T-cells were gated as CD45+/CD19-/CD3+/CD4+. Asterisks denote the significance between either F-CH and F-HF mice or M-CH and M-HF mice (* $p < 0.05$, *** $p < 0.001$, ns = not significant). Unpaired Mann-Whitney U t-test was applied for testing. **(C)** Percentage of CD8+ T-cells found in the peripheral blood of F-CH, F-HF, M-CH, and M-HF diet study mice. CD8+ T-cells were gated as CD45+/CD19-/CD3+/CD8+. Asterisks denote the significance between either F-CH and F-HF mice or M-CH and M-HF mice (* $p < 0.05$, ** $p < 0.01$, ns = not significant)**.** Unpaired Mann-Whitney U t-test was applied for testing. F-CH, female chow; F-HF, female high-fat; M-CH, male chow; M-HF, male high-fat.

**Supplementary Figure S4**

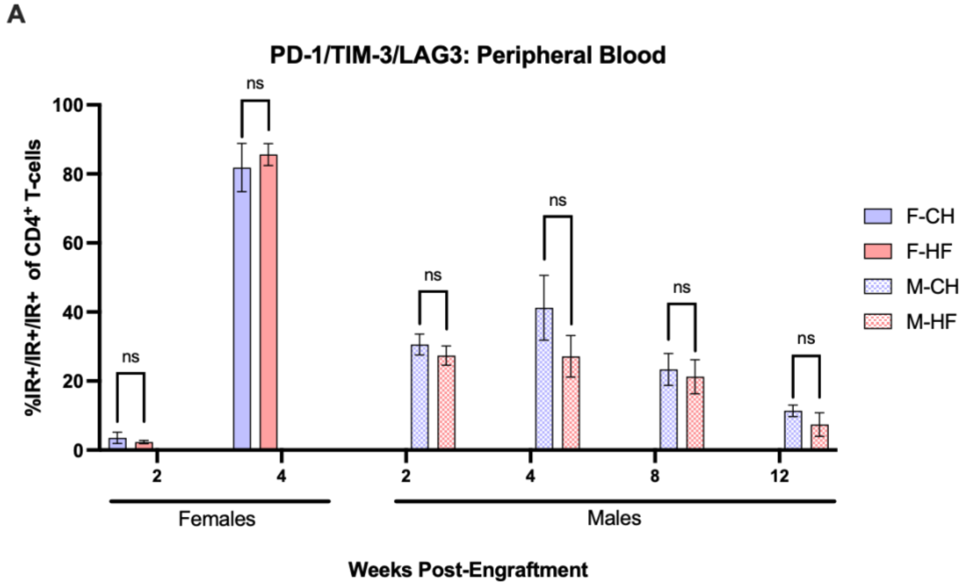


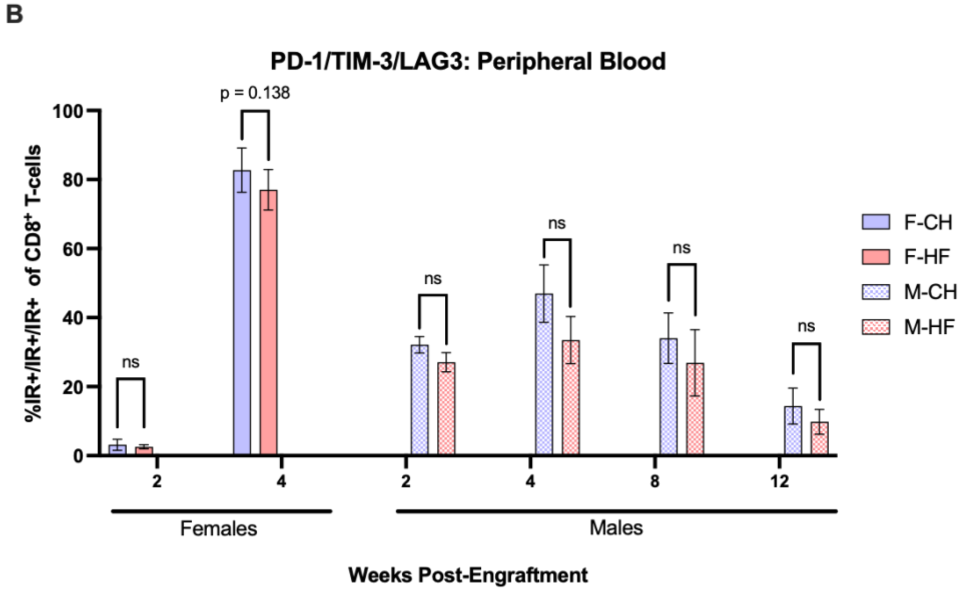


**Supplementary Figure S4. Expression of inhibitory receptors on T-cell subsets in the peripheral blood of female and male diet study mice. (A)** Percentage of peripheral blood CD4+ T-cells co-expressing immune inhibitory receptors PD-1, TIM-3, and LAG3 in diet study mice (F-CH, n = 10; F-HF, n = 10; M-CH, n = 10; M-HF, n = 10). Cells were gated as CD45+/CD19-/CD3+/CD4+/PD-1+/TIM-3+/LAG3+. Unpaired Mann-Whitney U t-test was applied for testing (ns = not significant). **(B)** Percentage of peripheral blood CD8+ T-cells co-expressing immune inhibitory receptors PD-1, TIM-3, and LAG3 in diet study mice. Cells were gated as CD45+/CD19-/CD3+/CD8+/PD-1+/TIM-3+/LAG3+. Unpaired Mann-Whitney U t-test was applied for testing (ns = not significant). F-CH, female chow; F-HF, female high-fat; M-CH, male chow; M-HF, male high-fat.

#### Supplementary Figure S5

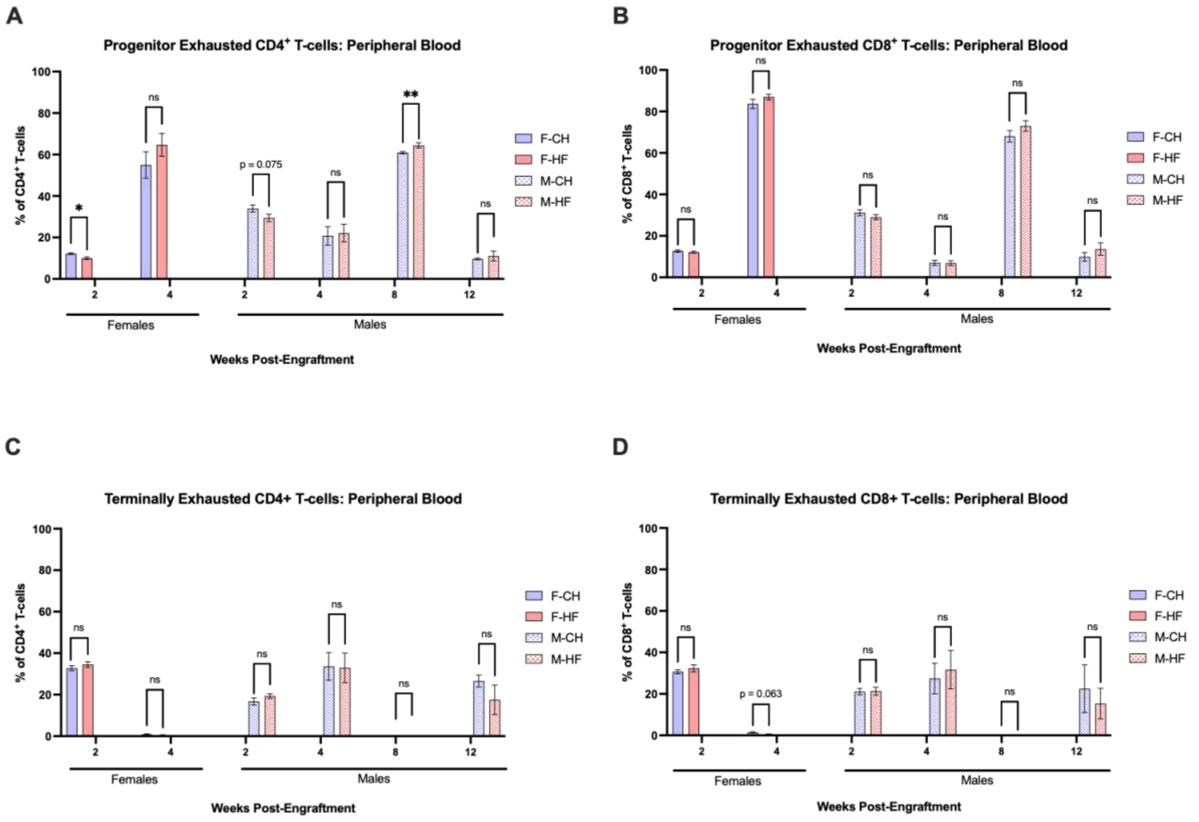


**Supplementary Figure S5. Characterization of exhaustion on T-cell subsets in the peripheral blood of female and male diet study mice. (A)** Percentage of peripheral blood CD4+ T-cells categorized into progenitor exhausted (TPEX; PD-$1^{int}$/TIM-$3^{lo/-}$) T-cells. Asterisks denote the significance between F-CH and F-HF or M-CH and M-HF diet study mice (* $p < 0.05$, ** $p < 0.001$, ns = not significant). Unpaired Mann-Whitney U t-test was applied for testing. **(B)** Percentage of peripheral blood CD8+ T-cells categorized into progenitor exhausted (TPEX; PD-$1^{int}$/TIM-$3^{lo/-}$) T-cells. Unpaired Mann-Whitney U t-test was applied for testing (ns = not significant). **(C)** Percentage of peripheral blood CD4+ T-cells categorized into terminally exhausted (TTEX; PD-$1^{hi}$/TIM-$3^{hi}$) T-cells. Unpaired Mann-Whitney U t-test was applied for testing (ns = not significant). **(D)** Percentage of peripheral blood CD8+ T-cells categorized into terminally exhausted (TTEX; PD-$1^{hi}$/TIM-$3^{hi}$) T-cells. Unpaired Mann-Whitney U t-test was applied for testing (ns = not significant).

#### Supplementary Figure S6

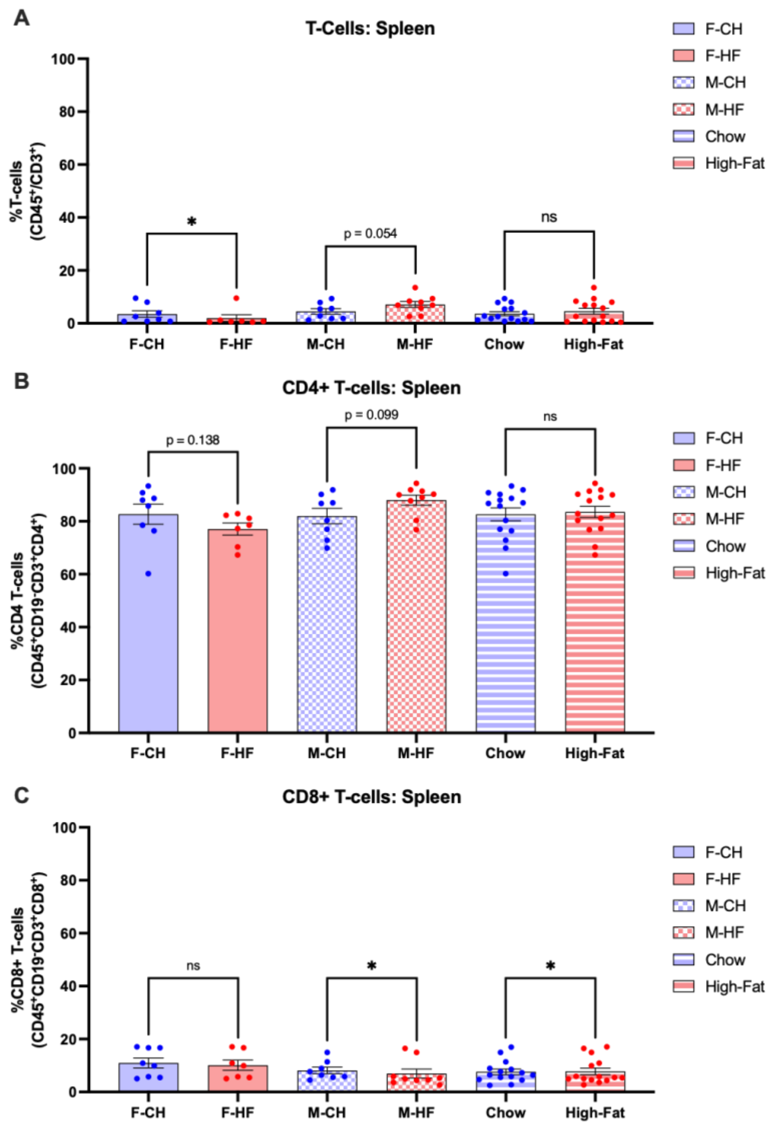


**Supplementary Figure S6. Characterization of T-cell subsets in the spleens of female and male diet study mice. (A)** Percentage of splenic T-cells in diet study mice (F-CH, n = 10; F-HF, n = 10; M-CH, n = 10; M-HF, n = 10). T-cells were gated as Live/CD45+/CD19-/CD3+. Asterisks denote the significance between either F-CH and F-HF mice or M-CH and M-HF mice (* $p < 0.05$, ns = not significant). Unpaired Mann-Whitney U t-test was applied for testing. **(B)** Percentage of splenic CD4+ T-cells in F-CH, F-HF, M-CH, and M-HF diet study mice. CD4+ T-cells were gated as Live/CD45+/CD19-/CD3+/CD4+. Asterisks denote the significance between either F-CH and F-HF mice or M-CH and M-HF mice (ns = not significant). Unpaired Mann-Whitney U t-test was applied for testing. **(C)** Percentage of splenic CD8+ T-cells in F-CH, F-HF, M-CH, and M-HF diet study mice. CD8+ T-cells were gated as Live/CD45+/CD19-/CD3+/CD8+. Asterisks denote the significance between either F-CH and F-HF mice or M-CH and M-HF mice (* $p < 0.05$, ns = not significant**).** Unpaired Mann-Whitney U t-test was applied for testing. F-CH, female chow; F-HF, female high-fat; M-CH, male chow; M-HF, male high-fat.

**Supplementary Figure S7**

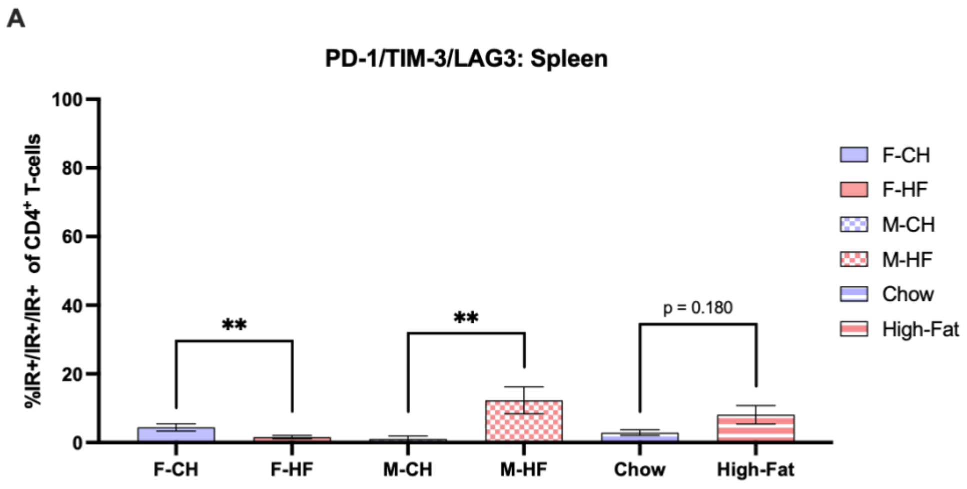


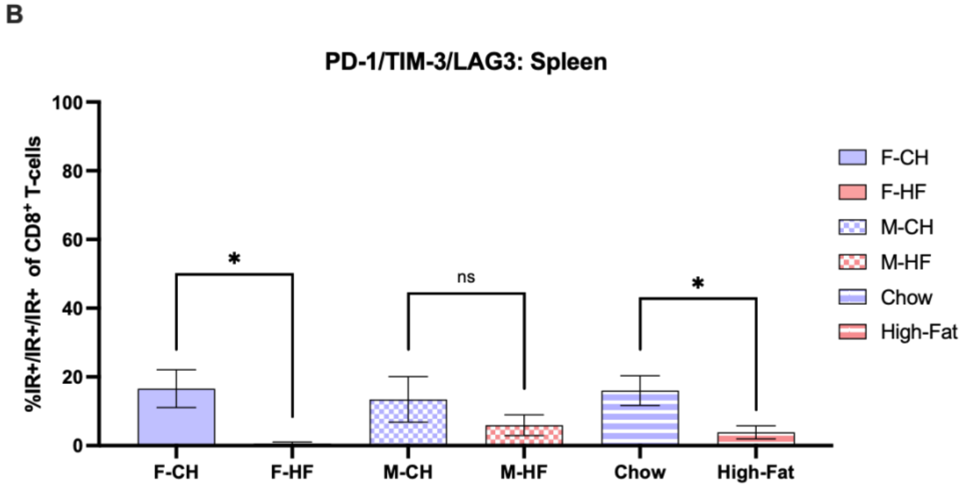


**Supplementary Figure S7. Expression of inhibitory receptors on splenic T-cell subsets of female and male diet study mice. (A)** Percentage of splenic CD4+ T-cells co-expressing immune inhibitory receptors PD-1, TIM-3, and LAG3 in diet study mice (F-CH, n = 10; F-HF, n = 10; M-CH, n = 10; M-HF, n = 10). Cells were gated as Live/CD45+/CD19-/CD3+/CD4+/PD-1+/TIM-3+/LAG3+. Asterisks denote the significance between F-CH and F-HF mice, M-CH and M-HF mice, or CH and HF mice (** $p < 0.001$). Unpaired Mann-Whitney U t-test was applied for testing. **(B)** Percentage of splenic CD8+ T-cells co-expressing immune inhibitory receptors PD-1, TIM-3, and LAG3 in diet study mice. Cells were gated as Live/CD45+/CD19-/CD3+/CD8+/PD-1+/TIM-3+/LAG3+. Asterisks denote the significance between F-CH and F-HF mice, M-CH and M-HF mice, or CH and HF mice (* $p < 0.01$). Unpaired Mann-Whitney U t-test was applied for testing. F-CH, female chow; F-HF, female high-fat; M-CH, male chow; M-HF, male high-fat.

**Supplementary Figure S8**

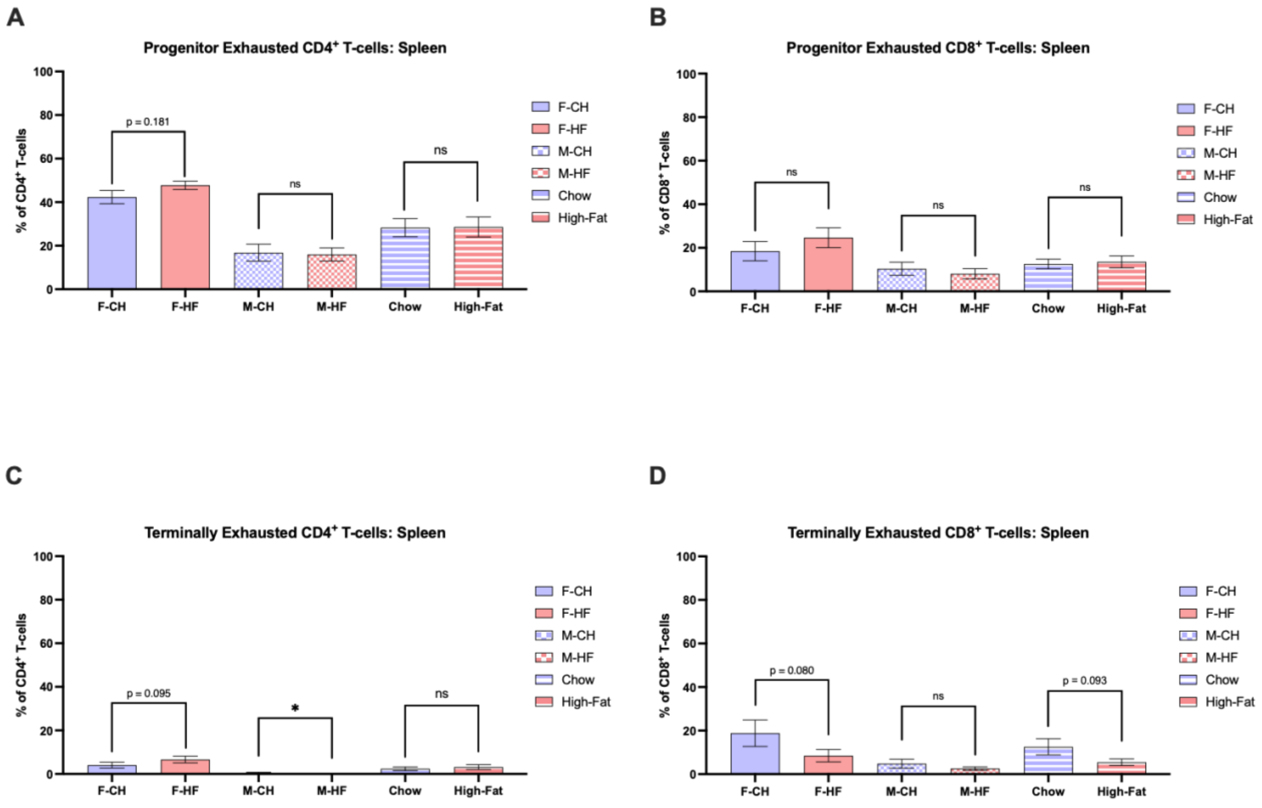


**Supplementary Figure S8. Characterization of exhaustion on splenic T-cells of female and male diet study mice. (A)** Percentage of splenic CD4+ T-cells categorized into progenitor exhausted (TPEX; PD-$1^{int}$/TIM-$3^{lo/-}$) T-cells. Unpaired Mann-Whitney U t-test was applied for testing (ns = not significant). **(B)** Percentage of splenic CD8+ T-cells categorized into progenitor exhausted (TPEX; PD-$1^{int}$/TIM-$3^{lo/-}$) T-cells. Unpaired Mann-Whitney U t-test was applied for testing (ns = not significant). **(C)** Percentage of peripheral blood CD4+ T-cells categorized into terminally exhausted (TTEX; PD-$1^{hi}$/TIM-$3^{hi}$) T-cells. Asterisks denote the significance between F-CH and F-HF, M-CH and M-HF, or CH and HF (* $p < 0.05$, ns = not significant). Unpaired Mann-Whitney U t-test was applied for testing. **(D)** Percentage of splenic CD8+ T-cells categorized into terminally exhausted (TTEX; PD-$1^{hi}$/TIM-$3^{hi}$) T-cells. Unpaired Mann-Whitney U t-test was applied for testing (ns = not significant).

#### 2.2 Supplementary Tables

**Supplementary Table 1.** Summary of study diets based on Teklad Diets' nutrient data sheets.

| | **Chow** [a] | **High-Fat** [b] |
|---|---|---|
| **Macronutrients (unit)** | | |
| *Protein (%)* | -- | 15.5 |
| *Crude Protein (%)* | 19.1 | -- |
| *Fat (ether extract; %)* [c] | 5.8 | -- |
| *Fat (%)* | -- | 16.7 |
| *Carbohydrate (available; %)* [d] | 44.3 | -- |
| *Carbohydrate (%)* | -- | 54.5 |
| *Fiber (%)* | -- | 3 |
| *Crude Fiber (%)* | 4.6 | -- |
| *Natural Detergent Fiber (%)* [e] | 13.7 | -- |
| *Calories from:* | | |
| *Protein (%)* | 25 | 15.5 |
| *Fat (%)* | 17 | 34.5 |
| *Carbohydrates (%)* | 58 | 50 |
| **Micronutrients: minerals (units)** | | |
| *Calcium (%; g/kg)* | 1 | 2 |
| *Chlorine (g/kg)* | -- | 10.8 |
| *Chloride (%)* | 0.5 | -- |
| *Chromium (mg/kg)* | -- | 1 |
| *Copper (mg/kg)* | 23 | 2.8 |
| *Iodine (mg/kg)* | 3 | 0.21 |
| *Iron (mg/kg)* | 240 | 31.9 |
| *Magnesium (%; g/kg)* | 0.2 | 0.588 |
| *Manganese (mg/kg)* | 93 | 10.5 |
| *Molybdenum (mg/kg)* | -- | 0.15 |
| *Phosphorus (%; g/kg)* | 0.7 | 2.7 |
| *Potassium (%; g/kg)* | 0.8 | 5.2 |
| *Selenium (mg/kg)* | 0.16 | 0.21 |
| *Sodium (%; g/kg)* | 0.3 | 7 |
| *Zinc (mg/kg)* | 63 | 30.3 |
| **Micronutrients: vitamins (unit)** | | |
| *Biotin (mg/kg)* | 0.77 | 0.2 |
| *Choline (mg/kg)* | 2200 | 642.6 |
| *Folate (mg/kg)* | 7 | -- |

| | | |
|---|---|---|
| *Folic Acid (mg/kg)* | -- | 1.3 |
| *Niacin (mg/kg)* | 100 | 50.7 |
| *Pantothenate (mg/kg)* | -- | 14.7 |
| *Pantothenic Acid (mg/kg)* | 87 | -- |
| *Vitamin A (IU/g; IU/kg)* | 30 | 4318 |
| *Vitamin B1 (thiamin; mg/kg)* | 95 | 3.2 |
| *Vitamin B12 (cyanocobalamin; mg/kg)* | 0.09 | 0.01 |
| *Vitamin B2 (riboflavin; mg/kg)* | 14 | 4.4 |
| *Vitamin B6 (pyridoxine; mg/kg)* | 17 | 4 |
| *Vitamin C (mg/kg)* | -- | 0 |
| *Vitamin D (IU/g; IU/kg)* | 2.4 | 400 |
| *Vitamin E (IU/kg)* | 150 | 25 |
| *Vitamin K (mg/kg)* | -- | 0.2 |
| *Vitamin K3 (menadione; mg/kg)* | 80 | -- |

For both diets, values are calculated from data of included ingredients. Actual values of any given batch of diet may vary slightly.
[a] Chow diet refers to the following Teklad Diet: 7012 – LM-485 Sterilizable Mouse/Rat Diet. 7012 is a fixed formula diet designed to support growth and reproduction of rodents. 7012 is supplemented with additional vitamins to ensure nutritional adequacy after autoclaving.
[b] High-fat diet refers to the following Teklad Diet: TD.110424 – New Total Western Diet. TD.110424 is a Teklad™ Custom Diet designed based on customer specifications of nutrient values based on NHANES and the Total Western Diet mean data.
[c] Ether extract is used to measure fat in pelleted diets.
[d] Carbohydrate (available) is calculated by subtracting neutral detergent fiber from total carbohydrates.
[e] Neutral detergent fiber is an estimate of insoluble fiber, including cellulose, hemicellulose, and lignin. Crude fiber methodology underestimates total fiber.

**Supplementary Table 2.** Multivariable Cox proportional hazard model comparing the effects of gender and diet on days-to-event

| Model | Source | Parameter | | Reference | Estimate | P value | Hazard Ratio (95%Confidence Interval) |
|---|---|---|---|---|---|---|---|
| Multivariable model without interaction | Gender | Male | | Female | -1.687 | <0.001 | 0.185 ( 0.081 - 0.422 ) |
| | Diet | Chow | | High Fat | -1.221 | 0.002 | 0.295 ( 0.136 - 0.638 ) |
| | **Variable** | **Class** | | | **Estimate** | **P value** | |
| Multivariable model with interaction | Gender | Male | | | -3.4725 | <0.0001 | |
| | Diet | Chow | | | -2.9562 | <0.0001 | |
| | Gender*Diet | Male | Chow | | 2.6417 | 0.0020 | |
